# Is increasing the effective leg length of a human runner metabolically beneficial?

**DOI:** 10.1101/2025.01.03.631222

**Authors:** Montgomery Bertschy, Herlandt Lino, Laura Healey, Wouter Hoogkamer

## Abstract

The purpose of this study was to understand if footwear midsole thickness can decrease the metabolic cost of running by increasing the effective leg length of a runner. We also sought to understand the role of midsole compliance on these measures.

Participants (n = 16) ran on a treadmill at 14 km h^-1^ in four mass-matched shoe conditions: 30, 40, and 60 mm thickness of a firm midsole foam and 60 mm thickness of a compliant midsole foam. Over two testing sessions, we measured metabolic cost (two 5-minute trials in each condition) and biomechanical measures (one 2-minute trial in each condition).

Increasing thickness in the firm midsole conditions increased effective leg length during early, mid, and late stance (all p < 0.001). However, metabolic cost increased (p = 0.031). Changing material from firm to compliant decreased effective leg length during midstance (p < 0.001), but not during early or late stance, and decreased metabolic cost (p < 0.001).

Results suggest that increasing midsole thickness can increase the effective leg length of a runner, but this isolated effect leads to increasing in metabolic cost. On the other hand, increasing midsole compliance leads to reductions in metabolic cost, agreeing with previous literature.

We suggest that the performance benefits seen in advanced footwear technology with increased midsole thickness are likely due to increased capacity to return mechanical energy, not due increases in effective leg length.

**Summary Statement:** Increased midsole compliance, not increased effective leg length, is responsible for decreased metabolic cost in running.

## Introduction

Running shoes have been increasing in thickness since the introduction of advanced footwear technology (AFT) in 2016 (Joubert & Jones, 2022). AFT is defined as performance running shoes with a highly compliant and resilient midsole, embedded stiffening elements, and a geometry that has a greater midsole thickness and more pronounced rounded geometry than traditional running footwear (Frederick, 2022). Since the introduction of AFT, significant improvements in elite road racing performances in nearly every distance have occurred (Bermon et al., 2021; Mason et al., 2024; Senefeld et al., 2021). Such improvements have resulted in controversy over footwear technology, resulting in the regulation of midsole thickness by World Athletics in 2020. These regulations were made based on the idea that increasing the thickness of footwear can increase the effective leg length of the runner, improving their performance (Burns & Tam, 2020).

Across the animal kingdom, greater effective limb length is correlated with lower metabolic cost of transport (Pontzer, 2007). However, it is not well understood if the effective limb length within a species can be modified to gain energetically efficient locomotion. A potential method to increase effective leg length in humans is footwear (Barrons et al., 2023). Theoretically, by increasing leg length with footwear of high midsole thickness without adding additional physiologically costly tissue, the runner may increase their stride length with minimal additional metabolic cost, thereby reducing the metabolic cost of running. However, this may not be entirely beneficial, as increased stride length from a longer effective leg length may actually increase the sagittal moment about the hip joint and the associated metabolic demands on hip muscles.

On the other hand, midsole thickness may play a much different role in running metabolic cost. Midsoles in AFT are compliant, resulting in greater midsole deformation under the same load than traditional footwear, often needing greater thickness to prevent bottoming out the midsole (the phenomenon where the midsole stiffens up exponentially when the deformation reaches the starting thickness) (Hoogkamer et al., 2018; Kram, 2022; Shorten, 2024). These midsole materials also tend to be highly resilient, showing high energy return in mechanical testing, but the effect of midsole energy return on the runner is highly debated (Kram, 2022; Matijevich et al., 2023; Shorten, 2024). Several studies have shown that softer midsole materials tend to reduce running metabolic cost (Frederick et al., 1986; Rodrigo-Carranza et al., 2024; Worobets et al., 2014), but there is also evidence that more underfoot cushioning does not always lead to improvements in metabolic cost (Hoogkamer, 2020; Tung et al., 2014)

Since more compliant midsole materials will lead to greater midsole compression, it is reasonable to assume that a firmer midsole material will be more likely to extend effective leg length throughout stance. Therefore, if the metabolic cost of running is decreased by increasing the midsole thickness with a firm material, then increased effective leg length may be a mechanism for reducing running metabolic cost in AFT. However, if the metabolic cost of running is decreased by increasing the compliance of a thick midsole, then increased energy absorption (and potentially return) may be a mechanism for improved running metabolic cost in AFT. The purpose of this study is to test the isolated effects of midsole thickness on running metabolic cost and biomechanics. The first aim is to test the concept of increased effective leg length as a mechanism for reducing the metabolic cost of running. We hypothesize that increased midsole thickness will lead to a reduced metabolic cost while running. The second aim is to test how midsole compliance affects the role of midsole thickness on the metabolic cost of running. We hypothesize that greater midsole compliance will lead to a reduced metabolic cost while running. Additionally, we quantified joint mechanics to help provide future insights into the effects of midsole properties on running biomechanics and energetics.

## Methods

### Participants

The study protocol was approved by the University of Massachusetts institutional review board (IRB #2927). We recruited runners 18-45 years of age that were capable of running a sub-17:30 5-kilometer race or equivalent. Participants were free of musculoskeletal injury at the time of participation, and did not have surgery within 6 months of data collection. 18 (13M/5F) runners participated in the study. Two participants were unable to maintain a respiratory exchange ratio below 1.0 during metabolic testing and were removed from analysis. We analyzed data for the remaining 16 participants (12M/4F, Age: 25.4 ± 7.2, Height: 175.6 ± 8.8 cm, Mass: 65.3 ± 8.0 kg).

### Footwear conditions

We used five footwear conditions in this study: shoes with midsole thicknesses of 30 mm, 40 mm, and 60 mm, made from firm ethylene-vinyl acetate (EVA) foam (hardness: 52±2C), and shoes with a midsole thickness of 30 and 60 mm, made from a highly compliant polyether block amide (PEBA) based material (hardness: 43±2C). All footwear conditions were custom made based on the PUMA Deviate Nitro (30 mm compliant; see Figure 1) by adding additional layers of foam between the midsole and outsole, keeping embedded stiffening plate placement consistent relative to the top of the midsole. All footwear conditions were matched for weight based on the heaviest condition (60 mm firm, 465 g). Part 1 of the study included 30, 40, 60 mm firm, and 60 mm compliant conditions. Part 2 of the study included 30 mm firm, and 30 and 60 mm compliant conditions.

**Figure 1:**
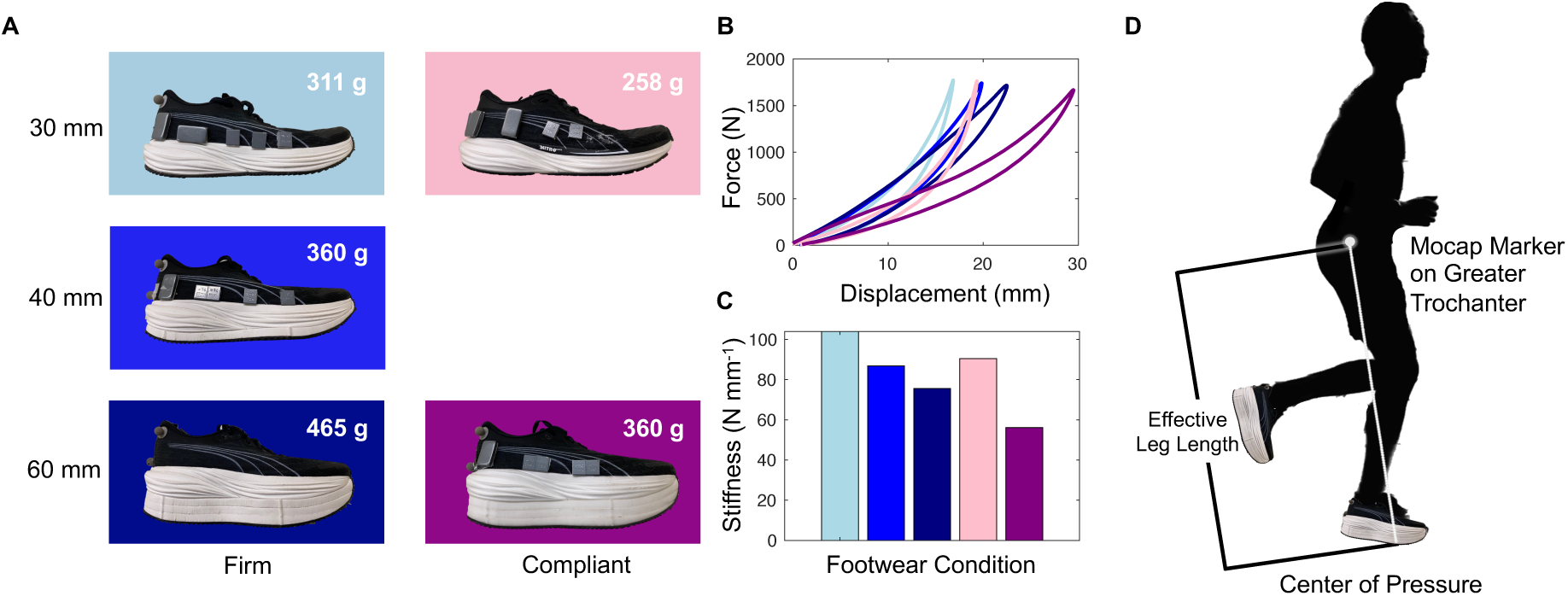
Summary of experimental footwear conditions. All footwear conditions were mass-matched to the 60 mm Firm condition (465 g) (A). 30 mm Compliant condition was used in an additional experiment presented in the Discussion. Colors used in plots B and C match the colors surrounding the respective footwear conditions in A. Effective leg length was calculated as the sagittal plane distance from the greater trochanter to the center of pressure (D).

### Mechanical Testing

The footwear compression stiffness, energy input, output, and percent return were measured using a material testing machine (Instron ElectroPuls 10000; Instron, Norwood, MA, USA). A rounded probe (diameter = 43 mm, radius of curvature = 21.5 mm) applied a parabolic force profile with a peak magnitude of 1800 N for 185 msec onto the heel of the shoe. Midsole compression stiffness was calculated from the maximum force applied and the maximum displacement (Table 1). Energy return was calculated as the ratio from the area under the force-deformation curve during unloading (output energy) to loading (input energy) (Table 1).

**Table 1:** Summary of mechanical testing results of footwear conditions.

| Thickness<br>(mm) | Material | Input<br>Energy (J) | Output<br>Energy (J) | Energy<br>Return (%) | Compression<br>Stiffness<br>(N mm<sup>-1</sup>) | Asker C<br>Hardness |
| --- | --- | --- | --- | --- | --- | --- |
| 30 | Firm | 9.8 | 7.0 | 71.9 | 104.0 | 52±2C |
| 40 | Firm | 13.7 | 9.9 | 72.1 | 86.8 | 52±2C |
| 60 | Firm | 17.3 | 12.3 | 71.1 | 75.6 | 52±2C |
| 30 | Compliant | 10.1 | 7.7 | 76.0 | 90.4 | 43±2C |
| 60 | Compliant | 20.5 | 15.3 | 74.5 | 56.2 | 43±2C |

**Table 2:** Summary of biomechanical measures. Values reported as mean ± s.d.

|  | Firm<br>30 mm | Firm<br>40 mm | Firm<br>60 mm | Compliant<br>60 mm | Thickness Effect |  | Material Effect |  |
| --- | --- | --- | --- | --- | --- | --- | --- | --- |
| | | | | | P-value | $\eta^2$ | P-value | $\eta^2$ |
| **Leg Length Early Stance**<br>**(cm)** | 94.2 ± 6.5 | 95.1 ± 6.3 | 97.0 ± 6.3 | 97.0 ± 6.1 | < 0.001 | 0.797 | 0.676 | 0.012 |
| **Leg Length Midstance**<br>**(cm)** | 86.9 ± 5.8 | 87.7 ± 5.9 | 89.5 ± 5.8 | 89.0 ± 5.8 | < 0.001 | 0.830 | < 0.001 | 0.745 |
| **Leg Length Late Stance**<br>**(cm)** | 94.6 ± 6.3 | 95.9 ± 6.4 | 97.3 ± 5.5 | 96.6 ± 6.3 | < 0.001 | 0.420 | 0.239 | 0.091 |
| **Leg Stiffness**<br>**(N mm<sup>-1</sup>)** | 24.4 ± 4.1 | 24.9 ± 6.9 | 23.6 ± 5.1 | 22.2 ± 4.8 | 0.398 | 0.023 | 0.006 | 0.405 |
| **Vertical Stiffness**<br>**(N mm<sup>-1</sup>)** | 37.1 ± 5.4 | 36.8 ± 5.9 | 36.2 ± 5.3 | 35.6 ± 5.3 | 0.018 | 0.166 | 0.178 | 0.117 |
| **Contact Time**<br>**(s)** | 0.217 ± 0.012 | 0.217 ± 0.012 | 0.218 ± 0.014 | 0.220 ± 0.013 | 0.159 | 0.063 | 0.038 | 0.258 |
| **Step length**<br>**(m)** | 1.33 ± 0.07 | 1.33 ± 0.07 | 1.33 ± 0.07 | 1.33 ± 0.07 | 0.594 | 0.009 | 0.693 | 0.011 |
| **Peak vGRF**<br>**(N)** | 1754 ± 276 | 1749 ± 262 | 1735 ± 272 | 1722 ± 260 | 0.085 | 0.093 | 0.169 | 0.122 |
| **Braking/Propulsive Impulse**<br>**(Ns)** | 43.8 ± 7.5 | 43.1 ± 7.1 | 42.5 ± 7.0 | 44.2 ± 7.3 | < 0.001 | 0.398 | < 0.001 | 0.748 |

### Experimental Protocol

Participants attended two data collection sessions: a metabolic session and a biomechanics session. The metabolic session consisted of a self-paced running warm up of at least 5 minutes in their own shoes. For familiarization participants ran 5 minutes in their own shoes on a force-instrumented, rigid treadmill (Treadmetrix, Park City, UT, USA) at the experimental speed of 14 km h^-1^ while breathing into a mouthpiece for indirect calorimetry. Participants then ran in each of the four shoe conditions (firm: 30, 40, 60 mm; compliant: 60 mm) twice in randomized, mirrored order (ABCDDCBA) for 5 minutes at 14 km h^-1^, followed by 5 minutes rest, for a total of eight running trials, while we measured expired air (TrueOne 2400, ParvoMedics, Salt Lake City, UT, USA). The expired gas analysis system was calibrated before the first (ABCD) and second (DCBA) halves to reduce long-term analyzer drift. The outcome measure for this session was metabolic cost of running calculated as metabolic power (W kg^-1^).

The biomechanics session consisted of a self-paced running warm up of at least 5 minutes. Next, we placed 21 retro-reflective markers on the shoe tip, 1^st^ and 5^th^ metatarsal head, medial and lateral malleolus, medial and lateral epicondyle, greater trochanter, and left and right anterior and posterior superior iliac spine. Additional marker clusters were placed on the heel, shank, and thigh. Participants ran in each shoe condition in randomized order for 2 minutes while we measured kinematics (Oqus, Qualisys, Goteborg, Sweden) and ground reaction forces (GRFs) (Treadmetrix, Park City, UT, USA). The outcome measures for this experiment were leg length during stance (m), total and vertical stiffness of the leg and midsole (N mm^-1^), GRFs (N), step length (m), and joint angles, moments, and powers (degrees, Nm, W kg^-1^). There was no minimum duration between the two data collection sessions, as fatigue was minimal (e.g., some participants completed the sessions directly after one another).

### Data Analysis

We calculated metabolic power based on the rates of oxygen uptake and carbon dioxide production over the final 2 minutes of each trial (Péronnet & Massicotte, 1991). GRF data were collected at 1000 Hz and lowpass filtered with a 2^nd^ order dual pass Butterworth filter at 15 Hz. Heel strike was defined as the time when vertical GRF increased above 30 N, midstance as the time of peak vertical GRF, and toe off as the time when vertical GRF decreased below 30 N. Step length was calculated as the time from heel strike to heel strike of the opposite foot multiplied by the treadmill velocity. Contact time was time from heel strike to toe off. Effective leg length during stance was determined as the distance from the greater trochanter to the center of pressure during early stance (10% of stance), midstance, and late stance (90% of stance) (excluding the first and last portion of stance to remove extreme values of center of pressure location related to noise during the aerial phase and filtering artefacts) (Hoogkamer et al., 2019; Willwacher et al., 2014). We defined leg stiffness as the sagittal center of mass (estimated as the average locations of the left and right anterior and posterior superior iliac spine) displacement from heel strike to midstance divided by the peak vertical GRF (Arampatzis et al., 1999). We defined vertical stiffness as the vertical center of mass displacement from heel strike to midstance divided by the peak vertical GRF. Joint mechanics were processed in Visual 3D (C-Motion, Germantown, MD, USA) based on a 6-degrees of freedom model.

### Statistical Analysis

We used linear mixed effect models (Wilkinson et al., 2023) in our analysis using the basic formulas of 𝑂𝑢𝑡𝑐𝑜𝑚𝑒 𝑚𝑒𝑎𝑠𝑢𝑟𝑒 ∼ 𝐹𝑖𝑥𝑒𝑑 𝑒𝑓𝑓𝑒𝑐𝑡 + (1 | 𝑃𝑎𝑟𝑡𝑖𝑐𝑖𝑝𝑎𝑛𝑡), where the fixed effect is either midsole thickness (30, 40, 60 mm in firm material) or material (firm and compliant at 60 mm). Effect sizes were calculated as partial eta squared (η^2^). All previously described outcome measures were tested across midsole thickness and material at relevant timepoints (early, mid, late stance for leg length and vertical/leg stiffness; peak for joint angle, moment, and power) We performed all statistical analyses in R (R software Version 2022.12.0.353; The R Core Team, Vienna, Austria).

### Artificial Intelligence

R/Python/Matlab code for data analysis and figures were created with some assistance from ChatGPT (OpenAI, 2024).

## Results

### Midsole Thickness

Increasing thickness of the firm midsole material led to an increased leg length during early stance (p < 0.001; η^2^ = 0.797), midstance (p < 0.001; η^2^ = 0.830), and late stance (p < 0.001; η^2^ = 0.420) (Fig 2A-C; Table 1). Vertical stiffness was reduced as midsole thickness increased (p = 0.018; η^2^ = 0.166), but leg stiffness was unchanged.

**Figure 2:**
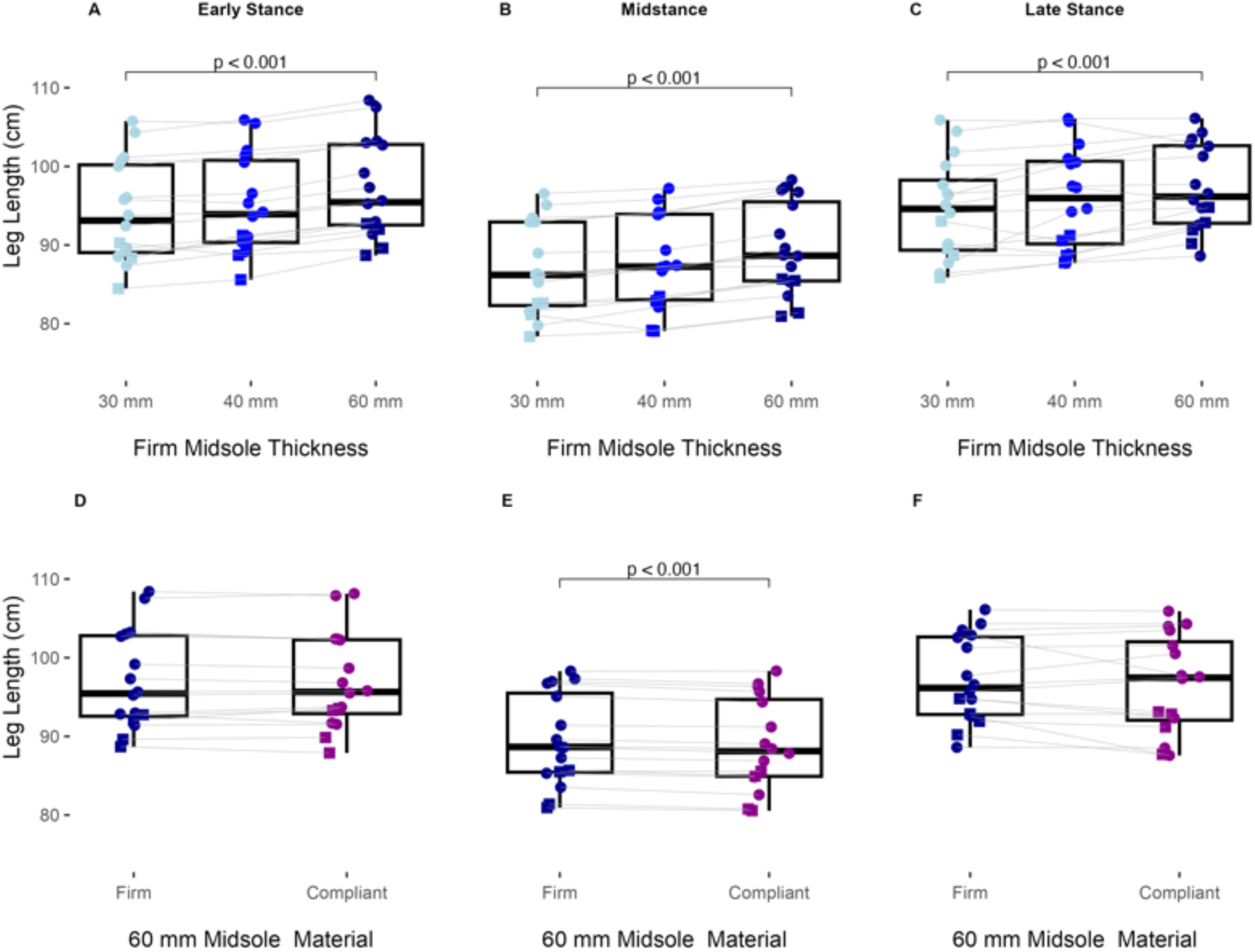
Leg length increases at heel strike, midstance, and toe off as midsole thickness increase; leg length decreases during midstance when midsole material is more compliant. Top row shows the effect of midsole thickness (A-C). Bottom row shows the effect of midsole material (D-F). Male and female participants are denoted by circles and squares, respectively.

Increasing thickness of a firm midsole material led to 0.4% increase in metabolic power per 10mm of added material (p = 0.031; η^2^ = 0.623) (Fig 3A). There was no effect of midsole thickness on step length, contact time, or peak vertical GRF. Braking and propulsive impulses decreased with increased midsole thickness (p < 0.001; η^2^ = 0.398).

**Figure 3:**
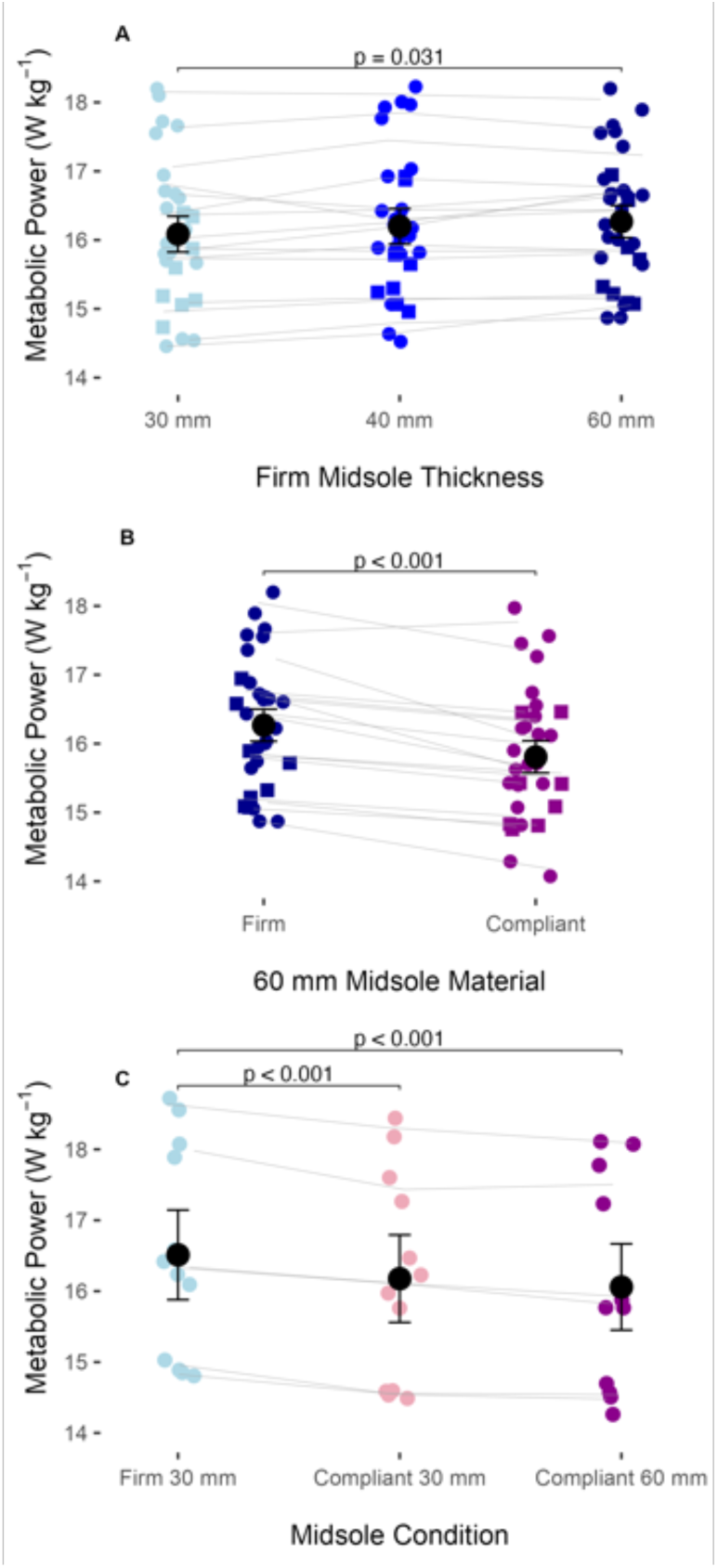
Metabolic power increased with increasing midsole thickness but decreased when midsole material was more compliant. For Part 1 (n = 16), the effect of midsole thickness (A) and midsole material (B) are presented. Male and female participants are denoted by circles and squares, respectively. For Part 2 (n = 6), all conditions are presented (C). Values are reported as mean ± s.e.m.

In the sagittal plane (Fig 4), increasing thickness of the firm midsole material led to a decrease in peak ankle dorsiflexion (p < 0.001; η^2^ = 0.547), plantarflexion moment (p < 0.001; η^2^ = 0.493), positive (p < 0.001; η^2^ = 0.652) and negative power (p < 0.001; η^2^ = 0.383), a decrease in peak knee flexion (p = 0.008; η^2^ = 0.139) and positive power (p = 0.002; η^2^ = 0.183), a decrease in positive hip moment (p = 0.020 ; η^2^ = 0.110), negative power (p = 0.038; η^2^ = 0.088), and an increase in hip positive power (p < 0.001; η^2^ = 0.227) at heel strike. In the frontal plane (Fig 5), increasing thickness of the firm midsole material led to an increase in peak ankle eversion (p < 0.001; η^2^ = 0.261), eversion moment (p = 0.028; η^2^ = 0.099), and positive (p = 0.033; η^2^ = 0.093) and negative power (p < 0.001; η^2^ = 0.295), but no differences at the knee or hip.

**Figure 4:**
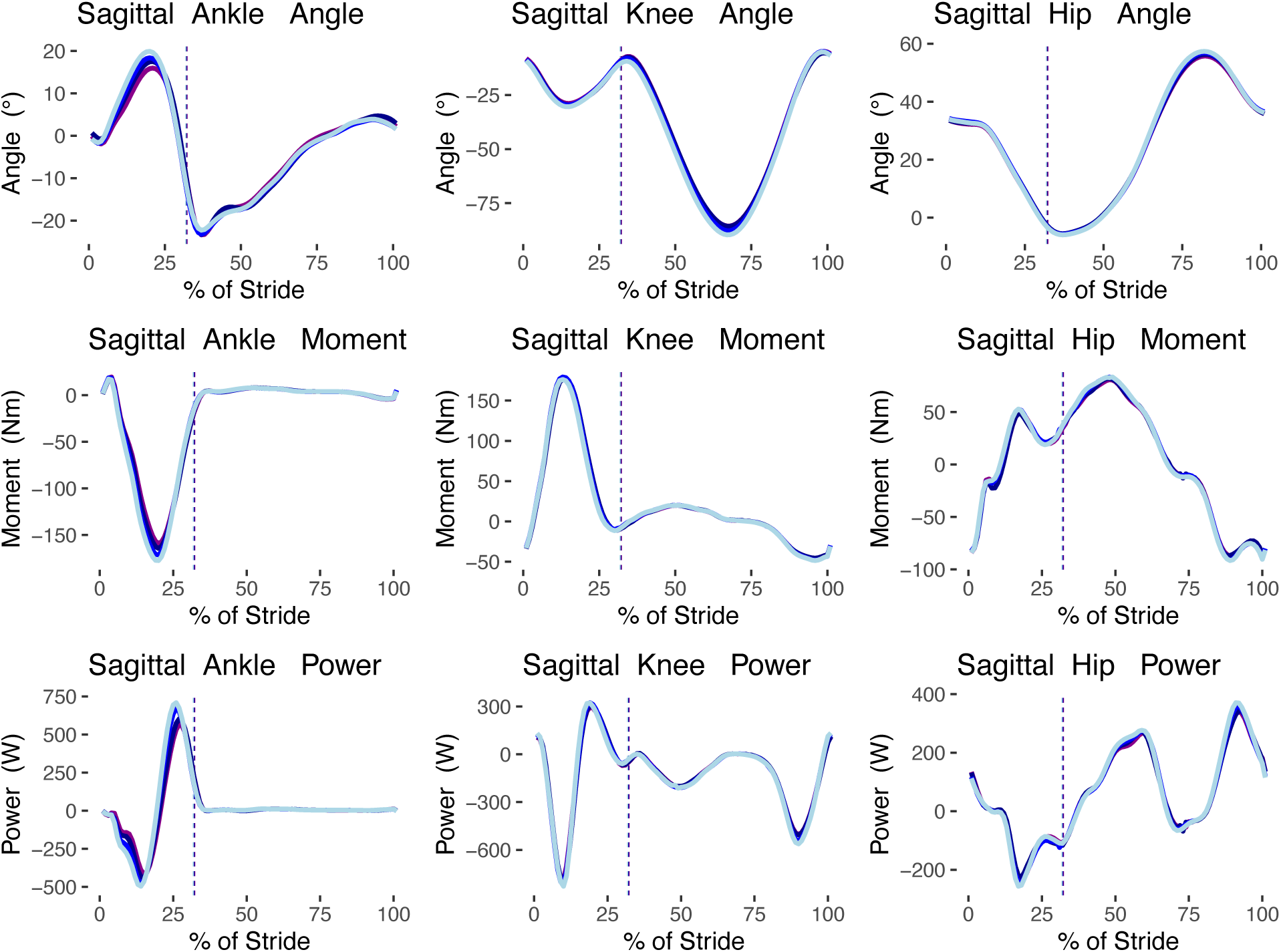
Summary of sagittal joint kinematics and kinetics.

**Figure 5:**
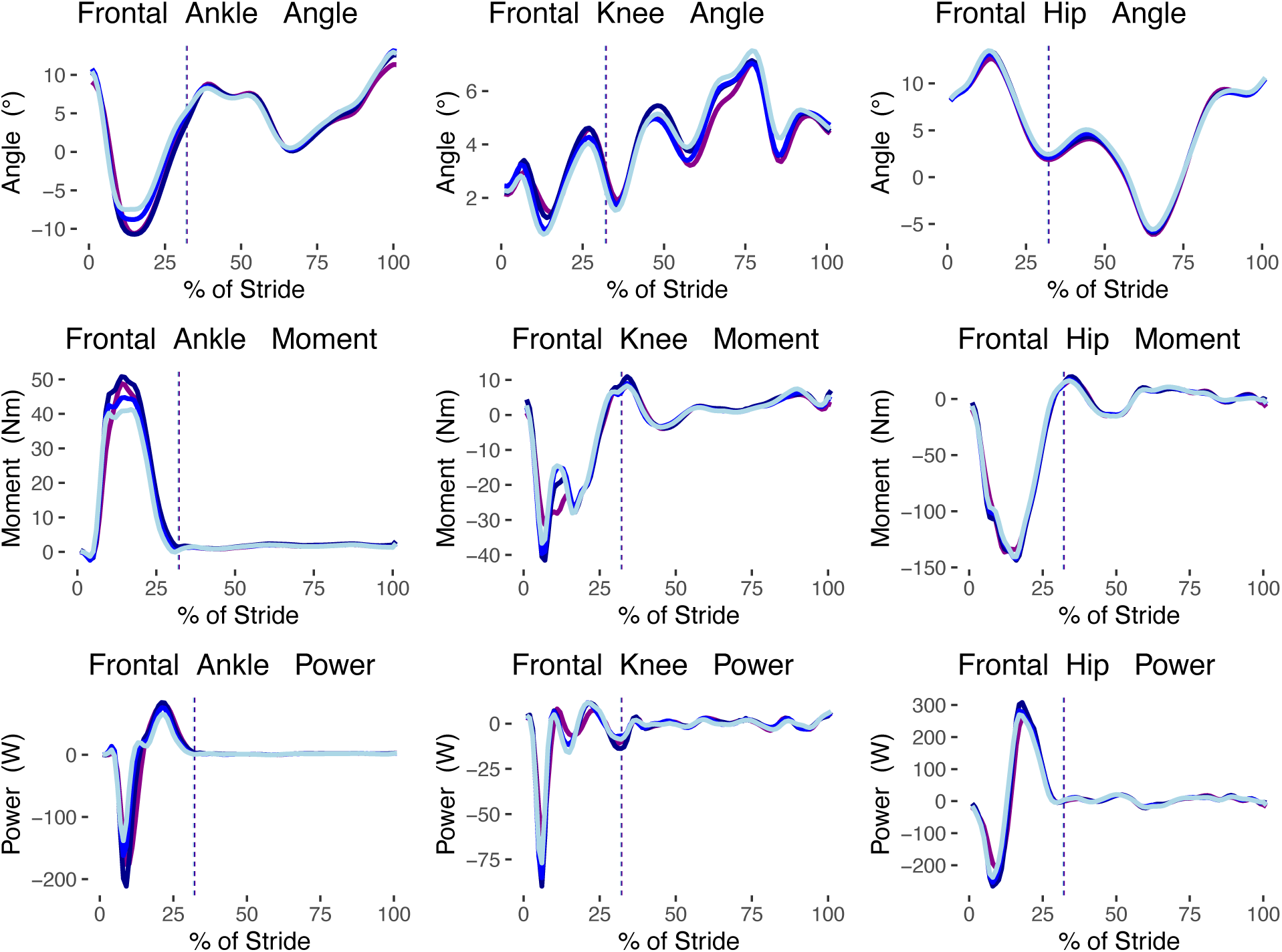
Summary of frontal joint kinematics and kinetics.

### Midsole Material

At the largest midsole thickness, the compliant material led to a significantly lower leg length at midstance (p < 0.001; η^2^ = 0.745), but not during early or late stance (Fig 2D-F; Table 1). Leg stiffness was significantly reduced from firm to compliant midsole material (p = 0.006; η^2^ = 0.405). Vertical stiffness was not different across midsole material.

At the 60 mm midsole thickness, the compliant material led to a 1.7% lower metabolic power than the firm material (p < 0.001; η^2^ = 0.387) (Fig 3B). Contact time increased from firm to compliant midsole material (p = 0.038; η^2^ = 0.258), but there were no significant changes in step length or peak vertical GRF. Braking and propulsive impulses increased firm to compliant material (p < 0.001; η^2^ = 0.748).

In the sagittal plane (Fig 4), at the largest midsole thickness, the compliant material had a significantly reduced peak ankle dorsiflexion (p = 0.037; η^2^ = 0.259), plantarflexion moment (p = 0.002; η^2^ = 0.471), and positive (p = 0.002; η^2^ = 0.485) and negative power (p = 0.0165; η^2^ = 0.327), but no differences in knee and hip mechanics. In the frontal plane (Fig 5), the compliant material had a significantly reduced peak ankle eversion moment (p = 0.012; η^2^ = 0.352) and negative power (p = 0.023; η^2^ = 0.301), reduced peak adduction knee moment (p = 0.004; η^2^ = 0.434) and negative power (0.006; η^2^ = 0.410), and reduced peak adduction hip angle (p < 0.001; η^2^ = 0.544), abduction moment (p = 0.010; η^2^ = 0.367), and positive (p = 0.001; η^2^ = 0.525) and negative power (p = 0.006; η^2^ = 0.411).

## Discussion

This study sought to determine if increasing the effective leg length of a runner reduces the metabolic cost of running. We found that effective leg length was increased at early, mid, and late stance as midsole thickness increased, achieving our goal for experimental conditions. This is in agreement with existing literature that has found midsole thickness to be related to effective leg length (Barrons et al., 2023). Despite running with an increased effective leg length, participants did not increase their step length. In overground running, step length can be increased while maintaining step frequency, resulting in an increased speed. Here participants ran at a fixed speed on a treadmill, so increasing step length would require a decrease in step frequency, which may not be advantageous for metabolic cost as it can result in suboptimal muscle-tendon dynamics (Swinnen et al., 2022). Yet, other studies have seen increases in step length in different footwear conditions that led to favorable metabolic outcomes ( Franz et al., 2012; Hoogkamer et al., 2018; Hunter et al., 2019; Rodrigo-Carranza et al., 2024).

Contrary to our hypothesis, as effective leg length increased, metabolic cost did not decrease, but increased. While the relationship between leg length and metabolic cost of transport is strong across the animal kingdom (Pontzer, 2007), this does not appear to be the case within humans.

These findings also diverge from the findings of Barrons et al., who reported no difference in metabolic cost across midsole thickness (Barrons et al., 2023). There are several distinctions to be made regarding the footwear conditions used in Barrons et al (2023) and our study. Firstly, the present study uses a larger range of footwear thickness (30 mm – 60 mm vs. 35 mm – 55 mm), which may explain why we do see a small, but significant, effect of thickness. Secondly, the footwear conditions in this study were purposely created with a firm EVA midsole material, whereas the footwear conditions used by Barrons et al. utilized a TPEE (thermoplastic polyester elastomer) that was substantially more compliant, which may play a role in how the thickness of the material may affect metabolic cost. It may also be that the effect of midsole thickness on metabolic cost is nonlinear, and the metabolically optimal thickness is at or below the thicknesses tested in these experiments. From minimal footwear to footwear with a midsole thickness of roughly 30 mm, larger effect of increasing midsole thickness on metabolic cost have been reported (Hébert-Losier et al., 2022; ), but those are likely related to other between shoe differences, such as midsole material and an embedded, stiffening plate. The findings of Tung et al. suggest that the isolated effect of cushioning (barefoot running on similar materials used in the present study) may be metabolically optimal at 10 mm of thickness, compared to 20 mm and no cushioning (Tung et al., 2014). However, Tung et al. purposely manipulated cushioning thickness without changing effective leg length, which was the purpose of the present study.

Braking and propulsive impulse decreased with increased midsole thickness, but there we no differences in contact time. While our purpose was to look at the effective leg length and energetics, we additionally quantified joint mechanics to help provide future insights into the effects of midsole properties on running biomechanics and energetics. We were primarily interested in evaluating if longer effective leg lengths would lead to longer step lengths, and if so, if that would lead to reduced metabolic rates, or if such longer steps would lead to increased hip extension moments in the sagittal plane offsetting the relative metabolic savings associated with longer steps. However, there was no difference in hip extension moment at heel strike.

Since there was no difference in step length, this is to be expected. Yet, there was a slight increase in positive power at the hip with increased midsole thickness at heel strike. Changes in metabolic cost of running are not easily explained by changes in joint mechanics, as there are many aspects that are not taken into account, such as power transfer from joint to joint, elastic storage and return form tendons, muscle fiber operating length and shortening velocity, different efficiencies for concentric and eccentric work, and many others (Beck et al., 2022; Bohm et al., 2023; Farris & Sawicki, 2011; Hata et al., 2024; Monte et al., 2020).

In the tallest condition (60 mm), the compliant midsole had a shorter effective leg length during midstance, but not during early and or late stance, compared to the firm midsole material. This confirms that the firm midsole resulted in a greater effective leg length during midstance than the compliant midsole. Additionally, this provides evidence that the compliant midsole indeed achieved greater midsole compression during running.

Further, the compliant midsole led to a decrease in metabolic cost compared to the firm midsole of the same thickness. This provides a strong indication that midsole material properties, not midsole thickness, accounts for the improvements in running performance seen in advanced footwear technology.

As midsole thickness increased in the firm material, the compression stiffness decreases. This is expected, as increasing thickness of the midsole acts as adding layers of midsole foam in series, where the total compression stiffness (k_eff_) is calculated as:

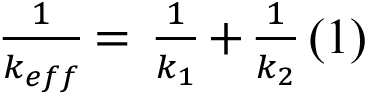

where k1 and k2 are the compression stiffness of each layer of midsole foam. Based on this, when the midsole thickness doubles from 30 mm to 60 mm, the midsole stiffness should be 50% lower. However, the stiffness only decreases from 104 N mm^-1^ to 75.6 N mm^-1^ (27.3% lower). This is due to the nonlinear stiffness of the foam materials, and the simplified method of calculating compression stiffness for this study. At a force of 1800 N (greater than what may occur at the heel during running), the lower midsole thickness approaches a “bottoming out” of the material, where the material no longer deforms under a force (see Force vs. Displacement graph in Fig 1). Additionally, the input and output energy increase, but percent energy return remains relatively unchanged. As expected, the shoe’s compression stiffness decreases when midsole material is changed from a firm to compliant material. The input and output energy increase, and percent energy return is also increased. The increase in percent energy return seems to be the only distinctive characteristic between midsole thickness effects and midsole material effects, suggesting that this is a factor in the savings in metabolic cost.

Vertical stiffness, from the center of mass to the running surface, decreased with increased midsole thickness. This decrease was small and is likely related to a combination of a larger increase in midsole compression and a small, insignificant decrease in peak vertical GRF. While the decrease in vertical stiffness is small, it may indicate that increased midsole compression led to larger center of mass displacements, rather than being counteracted by a stiffening of the leg to maintain center of mass mechanics as observed for altered surface condition (Daley & Biewener, 2006; Ferris et al., 1998, 1999). However, there was no effect of midsole thickness on two-dimensional leg stiffness. On the other hand, there was no significant effect of midsole material on vertical stiffness, but two-dimensional leg stiffness was decreased in the compliant material, likely in part due to the increase in braking impulse. This contrast in the effect of midsole thickness versus midsole material on vertical and leg stiffness highlights the difference in how the body adjusts to these footwear conditions and their material compression stiffness and energy return.

Sagittal plane kinetics suggest that increasing midsole thickness leads to large reductions in positive and negative power at the ankle, reductions in positive knee power, and reductions in negative hip power, yet a small to moderate increase in positive hip power at heel strike.

However, the frontal plane mechanics see increases in positive and negative ankle eversion power. While increased ankle eversion has not been directly linked to metabolic cost (Pizzuto et al., 2019), it is unlikely to be beneficial. Moreover, frontal plane mechanics are often discussed in relation to stability (Barrons et al., 2023), and increased midsole thickness has also been linked to increased whole body instability (Kettner et al., 2024 preprint). Based on these findings, the increase in metabolic power with increased midsole thickness may be related to frontal plane mechanics and instability, but further investigation is needed.

On the other hand, the compliant midsole led to a reduction positive (plantarflexion) and negative (dorsiflexion) power at the ankle, but no differences at the knee or hip in the sagittal plane. In the frontal plane, on the other hand, there were substantial decreases in joint powers at the ankle, knee, and hip in the compliant material. This contrasts with the increasing frontal plane ankle power due to midsole thickness. If increased midsole thickness leads to increased instability (Barrons et al., 2023; Kettner et al., 2024 preprint), it may be that increased midsole compliance may reduce the instability caused by increased midsole thickness, making the footwear better suited for long distance running.

The magnitude of the change in metabolic cost due to material (1.7% decrease from firm to compliant) is greater than the change due to midsole thickness (0.4% increase from per 10 mm added). For perspective, each 100 g added to the foot increases the metabolic cost of running by roughly 1% (Franz et al., 2012; Frederick et al., 1984). For these footwear conditions, the mass difference before equalizing was 154 g between the 30 mm and 60 mm firm conditions. This suggests that the estimated additional effect of added mass due to increased midsole thickness (∼1.54% increase in metabolic cost from 30 to 60 mm) would be larger than the effect due to increased thickness itself (1.14%). Therefore, when practically increasing midsole thickness (without mass-matching the baseline condition) the negative effect of added midsole thickness will likely be even greater than the results presented in this study.

Our initial findings provided us with an additional research question: if increasing the thickness of a firm midsole material leads to increased metabolic cost, does increasing the thickness of the compliant midsole change metabolic cost similarly? To answer this question, we set out to evaluate the metabolic cost of running in shoes with a 30 mm compliant midsole. We recruited 6 additional runners (6M, Age: 20.8 ± 3.1, Height: 176.0 ± 7.3 cm, Mass: 68.5 ± 5.5 kg) to look at the effect midsole thickness for the compliant material. Following the same warm up protocol as the metabolic session for the main experiment, participants ran in each of the three shoe conditions (Firm 30 mm; Compliant 30 and 60 mm) twice in randomized, mirrored order (ABCCBA) for 5 minutes at 14 km h^-1^, followed by 5 minutes rest, for a total of six running trials. Outcome measure for this session was metabolic power (W kg^-1^). At the 30 mm midsole thickness, the compliant material led to a 2.4% lower metabolic power than the firm material (p < 0.001; η^2^ = 0.538) (Fig 3C). However, we did not find a significant difference between the 30 mm and 60 mm compliant midsole conditions, nor was there a clear trend that could lead to speculation, which is to be expected with our relatively small sample size.

In this study, we looked at the effects of midsole thickness and midsole material on effective leg length and the metabolic cost of running. There are several measures that show similar trends between the two conditions. For example, peak ankle plantarflexion decreases as midsole thickness increases, just as peak ankle plantarflexion decreases from the firm to the compliant material. Similarly, midsole stiffness decreases as midsole thickness decreases and from the firm to the compliant material. However, there are some measures in which these two measures diverge. Specifically, metabolic power increases with increased midsole thickness, while metabolic power decreases from the firm to the compliant material. Additionally, frontal plane ankle mechanics tend to increase joint power for increased midsole thickness but decrease from the firm to the compliant material. Moreover, midsole percent energy return from mechanical testing increases from the firm to the compliant material but is mostly unchanged by midsole thickness. If runners are able to benefit from the energy returned from the midsole, this may by a mechanism for improved running metabolic cost in footwear with highly resilient materials (Matijevich et al., 2023).

### Limitations

This study has several limitations. Most notably, modifying one variable in footwear design will inherently affect another. For example, increasing midsole thickness will increase the longitudinal bending stiffness and the mass of the shoe. Regarding the former, all shoes in this study had a stiff plate, likely making the effect of increased stack height on longitudinal bending stiffness negligible between the conditions tested in Part 1, as they were all very stiff relative to on-the-market shoes. Additionally, the thicknesses of midsoles used in this study (30 mm to 60 mm) meant that all footwear conditions had a significant forefoot rocker that may not be present in thickness substantially less than 30 mm. Regarding mass, all shoes were mass-matched to the heaviest condition. As a result, however, all shoe conditions were far heavier that one could find in on-the-market running shoes. This also ignores the practical effects of adding midsole thickness to shoe. In reality, increased metabolic cost from adding midsole thickness would be accompanied by an increase in metabolic cost from added mass of the added thickness. Lastly, these footwear conditions may have a different effect if the participants were able to adapt over the course of several days or weeks. However, due to the impractical nature of the footwear conditions (mainly due to the mass), a longitudinal study may not be relevant. Nonetheless, this study adds to our understanding of specific features of advanced footwear technology.

### Summary

This study tested the hypothesis that increasing the midsole thickness of a running shoe can lead to an increased effective leg length, thereby decreasing the metabolic cost of running. Despite finding that effective leg length did indeed increase, metabolic cost also increased. On the other hand, our hypothesis that increased midsole compliance would reduce the metabolic cost of running was shown to be correct. We interpret this to indicate that the benefit of increased midsole thickness seen in advanced footwear technology is not due to increased effective leg length, but due to increased capacity for mechanical energy return. However, whether this mechanical energy can be returned to the runner has yet to be demonstrated directly.

## Acknowledgements

We thank John Kuzmeski and Zachary Barrons for mechanical testing of the footwear conditions.

## Data and resource availability

All relevant data and resource can be found within the article.

